# Acute stimulation of glucose metabolism by H₂O₂ sustains the NADPH steady-state under oxidative stress

**DOI:** 10.1101/2025.06.20.660777

**Authors:** C. Aburto, I. Ruminot, A. San Martín

## Abstract

Oxidative stress reprograms metabolic flux from glycolysis to the pentose phosphate pathway. Recently, it has been proposed that NADPH acts as a key molecule in pentose phosphate pathway regulation by exerting negative feedback through tonic inhibition of glucose-6-phosphate dehydrogenase. Interestingly, recent studies show that NADPH levels remain stable during acute exposure to hydrogen peroxide in the presence of glucose, ruling out NADPH-dependent feedback inhibition. We hypothesize that hydrogen peroxide triggers a feedforward activation mechanism, increasing NADPH production even before any detectable NADPH depletion. To probe this hypothesis, we used a panel of genetically encoded fluorescent indicators to monitor glucose, NADPH, F1,6BP, and pyruvate in single cells with high temporal resolution. Our results reveal that hydrogen peroxide rapidly activates glucose transport and consumption rates, enabling cells to preserve NADPH steady-state levels during early oxidative stress. Notably, this response precedes NADPH depletion, implying an anticipatory phenomenon that boosts NADPH production prior to its consumption. Furthermore, hydrogen peroxide induced an acute perturbation of F1,6BP steady-state and an increase of pyruvate accumulation. The pharmacological inhibition of the PPP’s gateway enzymes, glucose-6-phosphate dehydrogenase and transketolase, abolished the hydrogen peroxide-dependent alterations in F1,6BP steady-state levels and pyruvate accumulation, respectively. These findings suggest that a substantial fraction of glucose-derived carbon flux is diverted to the pentose phosphate pathway under oxidative stress, underscoring the importance of feedforward control in maintaining redox balance.

## INTRODUCTION

Oxidative stress induces a rerouting of metabolic flux from glycolysis to the pentose phosphate pathway (PPP) [1]. One proposed mechanism involves negative feedback via tonic inhibition of glucose-6-phosphate dehydrogenase (G6PDH) by NADPH [2]. However, recent evidence shows that NADPH levels do not decrease within the first seconds after acute hydrogen peroxide (H₂O₂) treatment in the presence of glucose, ruling out NADPH-dependent negative feedback inhibition [3]. A plausible alternative explanation is that H₂O₂ triggers a feedforward activation phenomenon, enhancing NADPH production even before NADPH depletion.

Previous studies indicate that prolonged H₂O₂ exposure can increase glucose transport activity [4, 5]. These findings are consistent with our observation that glucose is required to sustain NADPH levels under oxidative stress [3]. Therefore, the activation of glucose metabolism, together with an acute rerouting from glycolysis to the PPP, may be necessary to increase NADPH production and prevent its depletion.

In this study, we investigated whether H₂O₂ induces acute modulation of glucose metabolism and whether such modulation can sustain, in anticipatory manner, the steady-state NADPH levels during acute oxidative stress. In addition, we examined the extent to which glycolysis is acutely rerouted to the PPP, determining whether metabolic flux is preferentially directed toward the PPP. To address these questions, we employed a panel of Genetically Encoded Fluorescent Indicators (GEFIs) for glucose, NADPH, F1,6BP and pyruvate, which enabled us to investigate these processes at high temporal resolution at single-cell level.

## MATERIALS

NaCl, KCl, CaCl_2_, MgSO_4_ and HEPES were obtained from Sigma (St Louis, MO, USA). H₂O₂ (CAS N°107209) was purchased from Merck Millipore and a stock solution (1 M) prepared in water. D-(+)-Glucose (CAS N°50-99-7), Sodium L-Lactate (CAS N° 867-56-1) and Sodium Pyruvate (CAS N° 113-24-6) were purchased from Sigma-Aldrich and a stock solution (1 M) prepared in water stored at 4°C. G6PDi-1 (CAS N°2457232-14-1) was purchased from Sigma-Aldrich and prepared in DMSO stock (7 mM). Oxythiamine (CAS N° 136-16-3) was purchased from MedChem and a stock solution (100 mM) prepared in DMSO.

## METHODS

### Cell Culture and Transfection

HEK293 cells were acquired from the American Type Culture Collection (ATCC) and cultured at 37°C in 95% air/5% CO_2_ in DMEM/F12 10% fetal bovine serum. Cells were transfected at 60% confluency using Lipofectamine 3000 (Gibco) with 1 µg of each plasmid DNA obtained from Addgene: FLII12Pglu-700μΔ6/pcDNA3.1(-) (plasmid #17866), RedGlifon300/pcDNA3.1(-) (plasmid #163115), Pyronic/pcDNA3.1 (plasmid #51308), HYlight/pcDNA3.1 (plasmid #51308) and iNap1/pcDNA3.1-Hygro(+). Then cells were incubated for 16 hours with an efficiency of >60% and imaged at room temperature (22−25°C).

### Fluorescence measurements

For cell imaging experiments, cells were superfused with a solution of the following composition (in mM): 136 NaCl, 3 KCl, 1.25 CaCl_2_, 1.25 MgSO_4_, 10 HEPES pH 7.4, using an upright Olympus FV1000 confocal microscope equipped with a 20x water immersion objective (N.A. 1.0). 440, 488 and 543 nm solid-state lasers were used. Time series images were taken every one or two seconds with 20x (NA 1.0) in XYT scan mode (scan speed: 125 kHz; 256 × 256-pixel; pinhole 800 μm).

The fluorescent signal for both glucose and pyruvate sensors was converted to a concentration as follow: for glucose, we used a one-point calibration by applying a nominal zero (removing any carbon source), the reported K_D_ value of 660 µM and 50% ΔR_max_ [6]. For pyruvate, we employed a two-point calibration by applying a nominal zero (removing any carbon source), a saturating pulse of 10 mM pyruvate, and the reported K_D_ value of 107 µM [7]. We then substituted these data into the following equation to calculate the concentration of each metabolite: (*concentration*) = (*KD* ∗ *R*)/(*ΔRmax* − *R*), where R is the fluorescence ratio and Δmax is the maximum change in fluorescence.

### Data and statistical analysis

For data analysis, fluorescence signals were collected from regions of interest (ROIs) for each cell. Background subtraction was performed independently for each channel. The percentage change in fluorescence (%ΔF) was calculated from the difference between the maximum and minimum fluorescence values, using the formula: (*Fmax − Fmin*) ∗ *100*. Regression and statistical analysis were carried out using SigmaPlot (Jandel).

## RESULTS

### Hydrogen peroxide acutely activates glucose transport and consumption rates

H₂O₂ induces an acute increase in NADPH consumption rate [3]. However, NADPH steady-state levels remain unperturbed during the first seconds after the exposure to H₂O₂ in presence of glucose [3]. A plausible explanation for this phenomenon is that NADPH production rapidly increases due to simultaneous activation of glucose metabolism. To test this, we first explore if H₂O₂ induced a perturbation of steady-state levels of intracellular glucose in HEK293 cells. Indeed, a 30 second pulse of 250 µM H₂O₂ did not induce any change in the intracellular glucose concentration (**Figure 1A and B**). As intracellular glucose concentration does not inform about metabolic fluxes therefore, an alternative interpretation is that H₂O₂ induces a simultaneous activation in the rate of glucose consumption and uptake maintaining the intracellular glucose concentration invariant. Therefore, we decided to use two strategies to test glucose consumption and transport rates at high temporal resolution. First, we used the ITM (Inhibitor Transport Method) that uses the glucose transporter inhibitor cytochalasin B to perturb the glucose steady-state to dissect the glucose consumption rate [8]. Our results showed that H₂O₂ induced a rapid two-fold acceleration in the glucose consumption rate, increasing from 10.1 µM/s to 20.3 µM/s (**Figure 1C and D**). This result prompts us to assess the acute effect of H₂O₂ on glucose uptake. To do that, we evaluated the effect of H₂O₂ on glucose permeability as a readout of glucose transport. We used the 3-O-methyl-glucose (3OMG) method, which exploits trans-acceleration exchange. When 3-OMG is transported and released into the cytosol, the carrier’s inward-facing site becomes vacant, briefly increasing the likelihood that endogenous glucose binds and exits the cell. Simulations of glucose dynamics demonstrate that the initial rate of the 3-OMG-induced glucose decline is directly proportional to its permeability [9]. Therefore, enhanced glucose transport activity should appear as a steeper 3-OMG-induced decline in intracellular glucose concentration. Indeed, our results showed a clear H₂O₂-induced acceleration of glucose permeability observed as a faster decline in intracellular glucose concentration under 3-OMG pulse, indicating an acute increase in glucose transport of up to 3-fold, with a median change of 1.7-fold (**Figure 1E and F**). Therefore, the increase in glucose transport and consumption rate are similar due to peroxide treatment, explaining the lack of effect of H₂O₂ on the steady-state concentration of glucose. Altogether, these results indicate that H₂O₂ induces an acute, simultaneous activation of both glucose transport and consumption, thereby sustaining the PPP flux to maintain NADPH levels unchanged during the first seconds of exposure to oxidative stress.

**Figure 1.**
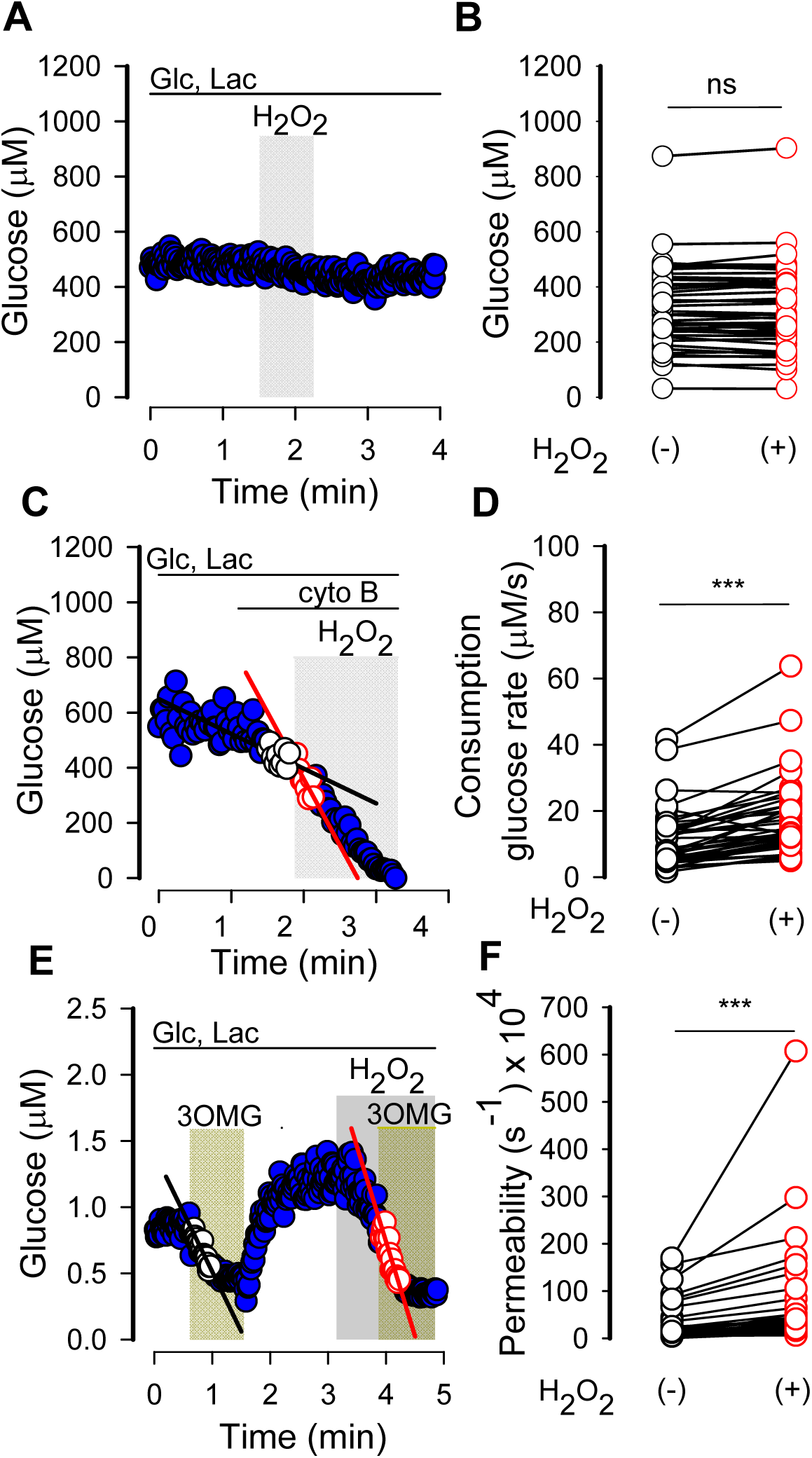
Hydrogen peroxide acutely activates glucose transport and consumption rates. **(A)** Representative trace of a HEK293 cell expressing glucose sensor perturbed with 250 µM H_2_O_2_ (grey shadow) in presence of 5 mM Glucose and 1 mM Lactate. Acquisition interval was 1 second. **(B)** Single cell before and after comparison of glucose to 250 µM H_2_O_2_. n= 45 cells from three independent experiments, ns: no significant p>0.05, Wilcoxon signed-rank test for paired samples was applied. **(C)** Representative trace response of glucose to 3OMG with and without H_2_O_2_. Circles with black or red edges and solid black or red lines represent the slopes used to measure glucose permeability in absence or presence of 250 µM H_2_O_2_, respectively. Acquisition interval was 1 second. **(D)** Single cell permeability quantification of glucose in absence and presence of H_2_O_2_. n=36 cells from three independent experiments, ***: p<0.001, Wilcoxon signed-rank test for paired samples was applied. **(E)** Representative trace response of glucose exposed to 20 µM cytochalasin B followed by a 250 µM H_2_O_2_ pulse in presence of 5 mM Glucose and 1 mM Lactate. Acquisition interval was 1 second. Circles with black or red edges and solid black or red lines represent the slopes used to measure consumption rates of glucose in absence or presence of 250 µM H_2_O_2_, respectively. **(F)** Single cell consumption glucose rate comparison before and after H_2_O_2_. n= 49 cells from five independent experiments, ***: p<0.001, Wilcoxon signed-rank test for paired samples was applied.

### Hydrogen peroxide-induced activation of glucose consumption precedes NADPH depletion

H₂O₂ induces an acute activation of glucose consumption rate. However, to sustain NADPH levels in high-demand conditions, the increase of glucose consumption must precede NADPH-depletion. Therefore, we monitored intracellular glucose and NADPH levels at single-cell resolution on a second-by-second basis. Our results show that the activation of glucose consumption precedes NADPH-depletion. Notably, NADPH consumption does not activate until eight seconds after H₂O₂ perturbation, whereas glucose consumption is activated within three seconds (**Figure 2A**). To determine whether anticipatory activation of the PPP sustains NADPH levels, we conducted a pharmacological loss-of-function experiment in which G6PDH was inhibited with 7 µM G6PDi-1 [10]. Indeed, the inhibition of the G6PDH completely abolished the delay time observed in the NADPH depletion and activation of glucose consumption rate synchronizing both phenomena under oxidative stress (**Figure 2B and 2C**). Also, the pharmacological inhibition of G6PDH increase the rate of activation of NADPH consumption under the treatment of H₂O₂ (**Figure 2D**). These findings indicate that H₂O₂-induced activation of glucose consumption indeed precedes NADPH depletion.

**Figure 2.**
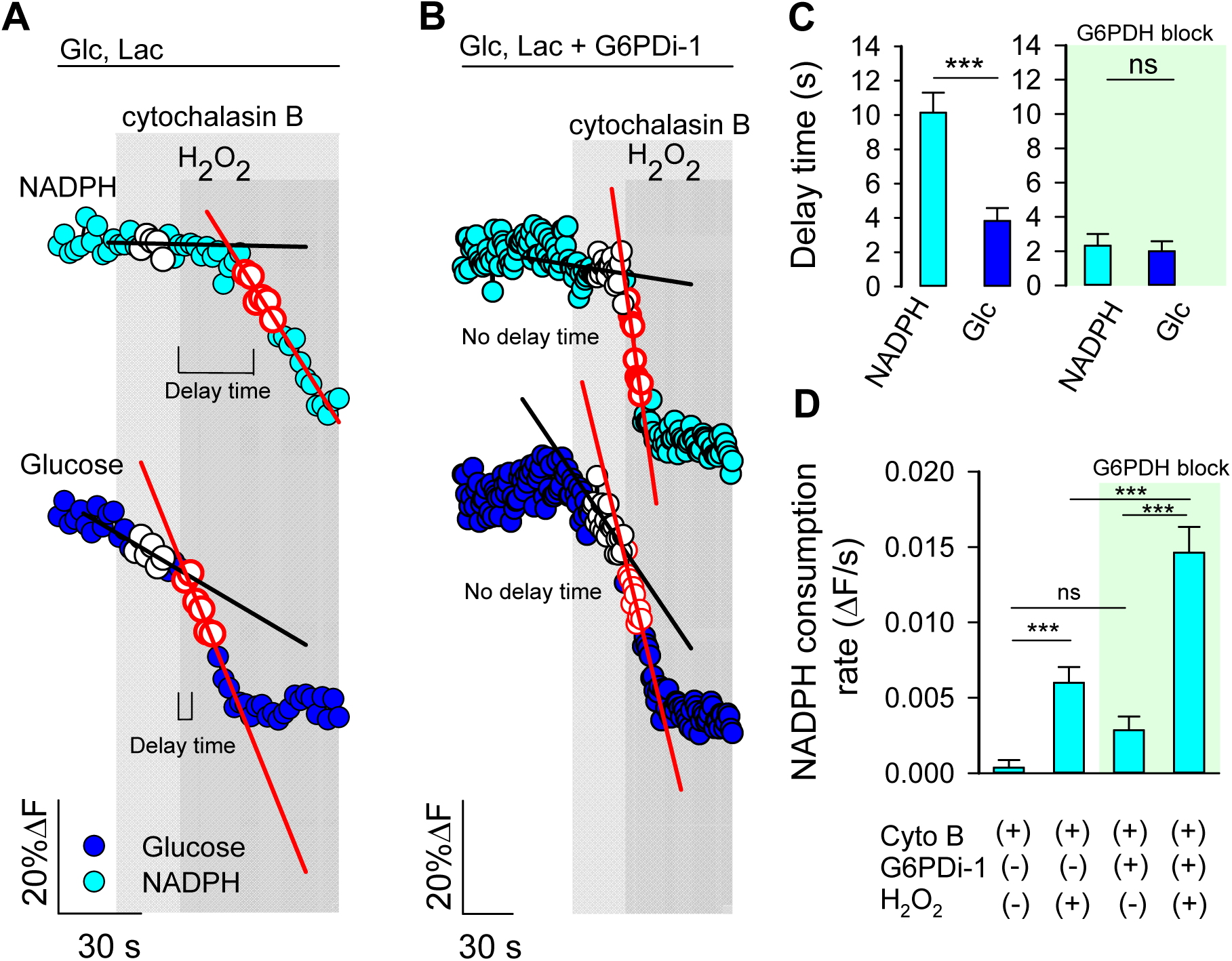
Hydrogen peroxide-induced activation of glucose consumption precedes NADPH depletion. **(A)** Representative trace of a HEK293 cell co-expressing iNap1 (cyan) and RedGlifon300 (blue) for NADPH and glucose, respectively. A 250 µM H_2_O_2_ pulse was applied on top of 20 µM cytochalasin B in the presence of 5 mM glucose, 1 mM lactate. Black-edged circles along with black lines represent the points taken for the rate measurement of each metabolite before H_2_O_2_, and red circles with red lines represent those under H_2_O_2_. Brackets (delay time) indicate the time between the addition of H_2_O_2_ and the break in the slope for NADPH and glucose. Acquisition interval was 1 second. **(B)** Same as (A), but in the presence of 7 µM G6PDi-1 to inhibit G6PD. No delay time was observed in this condition. **(C)** Comparison of delay times (in seconds) for iNap1 (NADPH) and RedGlifon300 (glucose) in response to cytochalasin B alone (left) or cytochalasin B plus G6PDi-1 (right, green panel). Data are shown as mean ± SEM from three independent experiments (17 cells for the left panel, 30 cells for the right panel). Paired Student’s t-test was performed. ns: p > 0.05, ***: p < 0.001. **(D)** Comparison of NADPH consumption rates before and after H_2_O_2_ stimulation, under cytochalasin B alone or cytochalasin B plus G6PDi-1 (green panel). Data are presented as mean ± SEM from three independent experiments. Statistical significance was assessed using paired Student’s t-test and Kruskal-Wallis test with Dunn’s post-test. ns: p > 0.05, **: p < 0.01, ***: p < 0.001.

### Hydrogen peroxide acutely activates PPP-derived pyruvate accumulation

H₂O₂ acutely activated glucose metabolism. This phenomenon reassembles the stimulation of glucose metabolism induced by neuronal signals such potassium [11] and nitric oxide [12, 13], which ultimately produced an activation of pyruvate accumulation. Therefore, we asked if this is the case for H₂O₂. Using a pharmacological pyruvate steady-state perturbation with the monocarboxylate transporter (MCT) inhibitor AR-C155858, we blocked the pyruvate efflux to assess its accumulation. Indeed, the MCT inhibition induces an intracellular pyruvate accumulation, which acutely accelerates under the exposure of H₂O₂ from 0.85 µM/s to 1.52 µM/s (**Figure 3A and B**). It has been described that H₂O₂ induced a two-fold increase of PPP flux [2]. In order to investigate the glycolytic or PPP source of the produced pyruvate, we blocked the non-oxidative phase of PPP using 10 µM of transketolase inhibitor oxythiamine. Surprisingly, this inhibition completely abolished the H₂O₂-induced pyruvate accumulation (**Figure 3C and D**), suggesting that a major fraction of the carbon flux from glucose is rerouted towards PPP under oxidative stress. Complementary to this data, we assessed the levels of fructose-1,6-bisphosphate (F1,6BP) using a recently developed single-fluorophore indicator HYlight [14]. This metabolite is located in the intersection of glycolysis and carbon re-injection from PPP. The exposure to H₂O₂ induced a decrease of steady-state of F1,6BP relative levels indicating a decrease of its production or an increase of its consumption (**Figure 3E**). However, the pharmacological inhibition of G6PDH with G6PDi-1 completely abolishes the effects of H₂O₂ over F1,6BP steady-state (**Figure 3E and F**), pointing out a decrease in its production. To obtain a semi-quantitative estimate of the sensor’s operating range, we ended every experiment with a maneuver that drove the sensors to their apo and/or saturated states (**Supplementary Figure 1**). Ours findings suggest that, under oxidative stress, a major fraction of the glucose-derived carbon flux is diverted to the PPP.

**Figure 3.**
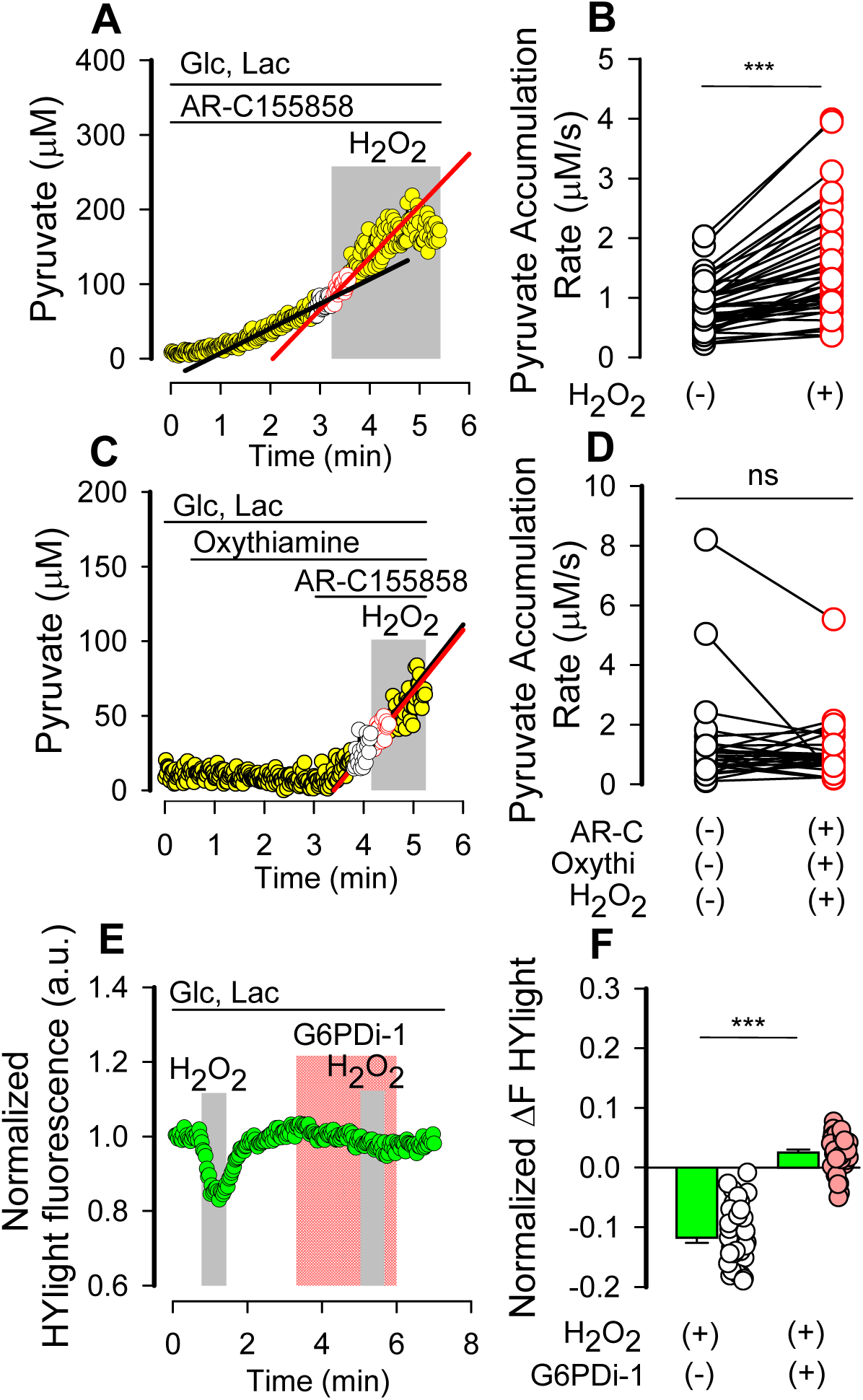
Hydrogen peroxide acutely activates pyruvate accumulation via non-oxPPP. **(A)** Representative trace of a HEK293 cell expressing a pyruvate sensor in which a 250 µM H_2_O_2_ pulse was applied in the presence of the MCT inhibitor. Black-edged circles along with black lines represent the points taken for the rate measurement of pyruvate before H_2_O_2_, and red circles with red lines represent those under H_2_O_2_. **(B)** Single-cell before and after comparison in the pyruvate production rate before and after the H_2_O_2_ pulse is shown. A median of 1.76 was observed. n = 40 cells, from four independent experiments, paired Student’s t-test was performed, ***: p < 0.001. **(C)** Representative trace of pyruvate in response to 250 µM H_2_O_2_ in the presence of 1 µM of the MCT inhibitor AR-C155858 and 10 µM oxythiamine. The slopes indicate the pyruvate accumulation rates before and after adding H_2_O_2_. Black-edged circles along with black lines represent the points taken for the rate measurement of pyruvate before H_2_O_2_, and red circles with red lines represent those under H_2_O_2_ **(D)** Single cell before and after comparison in the pyruvate production rate in response to 1 µM AR-C155858 and 10 µM oxythiamine in the presence or absence of 250 µM H_2_O_2_. n=39 cells from three independent experiments, paired Student’s t-test was performed. ns: no significant p>0.05. **(E)** Representative trace of a HEK293 cell expressing a F1,6BP sensor exposed to 250 µM H_2_O_2_ in presence and absence of 7 µM G6PDi-1. The acquisition interval was 2 seconds. **(F)** Comparison of F-1,6-BP ΔF values in response to 250 µM H_2_O_2_ in presence and absence of 7 µM G6PDi-1. Data are presented as the mean ± SE. n = 30 cells from three independent experiments, paired Student’s t-test was performed. ***: p < 0.001.

## DISCUSSION

Although oxidative stress induces an increase of NADPH consumption to replenish the reduced glutathione pool, acute H₂O₂ exposure does not disrupt NADPH steady-state concentration within the first seconds after treatment. In this study, we tested the hypothesis that H₂O₂ can acutely activate glucose metabolism, and that these activations precede the late drop in intracellular NADPH, thereby sustaining steady-state NADPH levels. Our results provide evidence of an acute activation of glucose transport and consumption rates, as far as we know, constitute the first report of acute H₂O₂ effects on glucose metabolism. These findings are consistent with earlier work reporting modulation of glucose transport, glycolysis and PPP activity stimulation after minutes to hours of H₂O₂ treatments [4, 5, 15–18].

Additionally, with the simultaneous measurements of glucose and NADPH at single-cell level, we were able to monitor these metabolites at high temporal resolution under oxidative stress. Notably, the activation of glucose consumption precedes NADPH-depletion by approximately six seconds, delay was completely abolished by the G6PDH inhibitor. This observation agrees with data from experiments in which G6PDH inhibition led to acute NADPH depletion during H₂O₂ exposure [3], indicating that glucose metabolism sustains NADPH steady-state in the initial seconds of oxidative stress. Based on the temporal profile of our findings, we hypothesize that the alternative mechanisms previously reported to divert glycolytic flux into the PPP [1, 2, 19] acts at a later time window, serving as a complementary response to sustained oxidative stress.

Interestingly, although our results showed that H₂O₂ induced an acute pyruvate accumulation, experiments using pharmacological inhibition of G6PDH and transketolase, gateways of PPP, suggest that an important fraction of the glycolytic flux is rerouted to PPP. Therefore, a substantial fraction of pyruvate does not arise from glycolysis but is instead generated through the PPP. These findings contribute to the ongoing debate regarding how representative isotope-based methods are in assessing the basal flux through the PPP under resting conditions, as well as the fraction of flux that shifts to the PPP upon activation [20]. However, our results were obtained from an immortalized cell line, which may not faithfully reflect the physiological properties of unaltered primary tissues or cells. For instance, phosphoglucose isomerase is a near-equilibrium enzyme that has been shown to be highly active at converting fructose-6-phosphate (F6P) into G6P allowing the re-cycling of carbon to the PPP in neurons, which is not observed in astrocytes where PPP-derived F6P is preferably converted to lactate [21]. Consequently, the fraction of G6P that re-enters the PPP versus that directed toward pyruvate formation is expected to vary among different cell types.

Our results further suggest that a major fraction of glycolytic flux is diverted to PPP during oxidative stress than previously described [2]. Although, in vivo studies in astrocytes shown that H₂O₂ stimulates glycolysis [15] our measurements of pyruvate-accumulation flux indicate that, whether glucose is catabolized through glycolysis or the PPP, its convergent end-product is pyruvate, a key glycolytic metabolite. In this regard, our data support the previous notion that oxidative stress triggers glycolytic activation. Because our assessment is semi quantitative, more precise studies using calibrated fluorescent indicators in a multiplexing measurement format at single cell resolution are needed, to determine exact flux values and dynamics through glycolytic and PPP pathways in resting and activated experimental conditions in different primary cells and tissues.

## CONCLUSIONS

This study provides compelling evidence that H₂O₂ acutely activates glucose metabolism, enabling cells to sustain NADPH steady-state levels during the first seconds of acute oxidative stress. Notably, this activation of glucose metabolism precedes NADPH depletion, supporting the existence of anticipatory phenomena that enhances NADPH production before its depletion. Pharmacological inhibition of the PPP’s gateway enzymes, G6PDH and transketolase, completely abolished both the H₂O₂-dependent steady-state F1,6BP perturbation and the activation of pyruvate accumulation, respectively. These findings suggest that a major fraction of glucose-derived carbon flux is rerouted to the PPP under oxidative stress.

## ACKNOWLEDGMENT

We thank the members of the Energy Metabolism Group at CECs and Felipe Barros for helpful discussions and support. We thank the website of app.Biorender.com for assistance in creating the graphical abstract (Created in BioRender. Aburto, C. (2025) https://BioRender.com/2003fss).

## AI USE STATEMENT

During the preparation of this work the author(s) used ChatGPT in order to improve language and readability. After using this tool/service, the author(s) reviewed and edited the content as needed and take(s) full responsibility for the content of the publication.

## SUPPLEMETARY FIGURE

**Supplementary figure 1.**
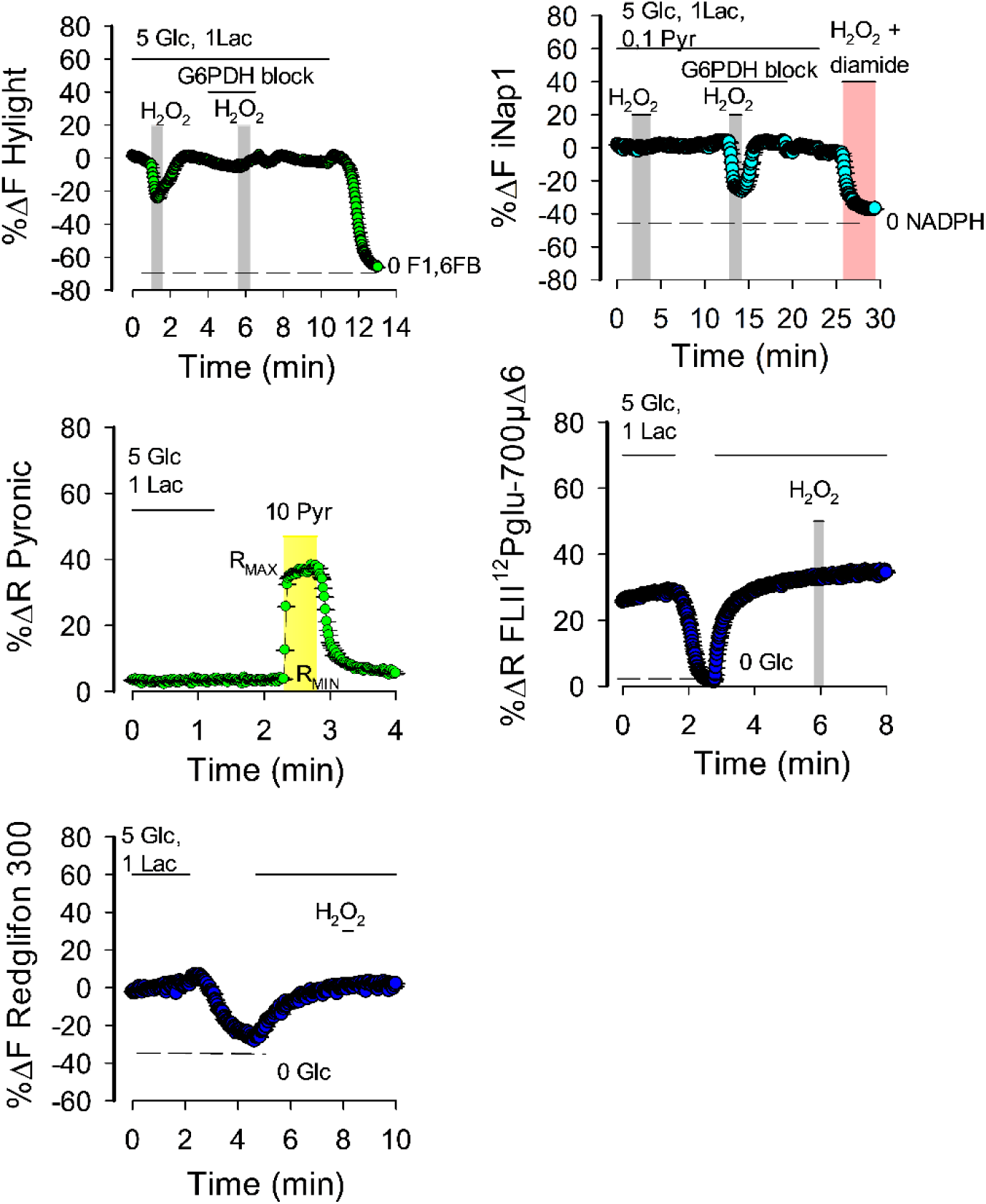
Control of fluorescent sensor behavior with F_MIN_ and F_MAX_ values under experimental conditions. **(A)** Representative experiment showing the minimum fluorescence of the Hylight sensor for F-1,6-BP upon removal of all carbon sources (5 mM glucose, 1 mM lactate). **(B)** Representative experiment showing the minimum fluorescence of the iNap1 sensor for NADPH following the addition of a potent oxidant pulse consisting of 1 mM H_2_O_2_ and 50 µM diamide. **(C)** Representative experiment showing the minimum and maximum fluorescence of the Pyronic sensor for pyruvate, achieved by removing all carbon sources (F_MIN_) and adding a saturating pulse of 10 mM pyruvate (F_MAX_). **(D)** Representative experiment showing the minimum fluorescence of the FLII^12^Pglu-700μΔ6 FRET sensor for glucose upon removal of all carbon sources (5 mM glucose, 1 mM lactate). **(E)** Representative experiment showing the minimum fluorescence of the RedGlifon300 SF sensor for glucose upon removal of all carbon sources (5 mM glucose, 1 mM lactate).

## REFERENCES

1. Ralser M, Wamelink MM, Kowald A, Gerisch B, Heeren G, Struys EA, et al. Dynamic rerouting of the carbohydrate flux is key to counteracting oxidative stress. J Biol. 2007;6(4):10. Epub 2007/12/25. doi: 10.1186/jbiol61. PubMed PMID: 18154684; PubMed Central PMCID: PMCPMC2373902.

2. Christodoulou D, Link H, Fuhrer T, Kochanowski K, Gerosa L, Sauer U. Reserve Flux Capacity in the Pentose Phosphate Pathway Enables Escherichia coli’s Rapid Response to Oxidative Stress. Cell Syst. 2018;6(5):569–78 e7. Epub 2018/05/14. doi: 10.1016/j.cels.2018.04.009. PubMed PMID: 29753645.

3. Aburto C, Parada-Goddard V, San Martín A. The activation of pentose phosphate pathway flux by hydrogen peroxide is not regulated by NADPH-mediated feedback inhibition. bioRxiv. 2025:2025.03.17.643767. doi: 10.1101/2025.03.17.643767.

4. Hamrahian AH, Zhang JZ, Elkhairi FS, Prasad R, Ismail-Beigi F. Activation of Glut1 glucose transporter in response to inhibition of oxidative phosphorylation. Arch Biochem Biophys. 1999;368(2):375–9. Epub 1999/08/12. doi: 10.1006/abbi.1999.1320. PubMed PMID: 10441390.

5. Prasad RK, Ismail-Beigi F. Mechanism of stimulation of glucose transport by H2O2: role of phospholipase C. Arch Biochem Biophys. 1999;362(1):113–22. Epub 1999/01/26. doi: 10.1006/abbi.1998.1026. PubMed PMID: 9917335.

6. Takanaga H, Chaudhuri B, Frommer WB. GLUT1 and GLUT9 as major contributors to glucose influx in HepG2 cells identified by a high sensitivity intramolecular FRET glucose sensor. Biochim Biophys Acta. 2008;1778(4):1091–9. Epub 2008/01/08. doi: 10.1016/j.bbamem.2007.11.015. PubMed PMID: 18177733; PubMed Central PMCID: PMCPMC2315637.

7. San Martin A, Ceballo S, Baeza-Lehnert F, Lerchundi R, Valdebenito R, Contreras-Baeza Y, et al. Imaging mitochondrial flux in single cells with a FRET sensor for pyruvate. PLoS One. 2014;9(1):e85780. Epub 2014/01/28. doi: 10.1371/journal.pone.0085780. PubMed PMID: 24465702; PubMed Central PMCID: PMCPMC3897509.

8. Bittner CX, Loaiza A, Ruminot I, Larenas V, Sotelo-Hitschfeld T, Gutierrez R, et al. High resolution measurement of the glycolytic rate. Front Neuroenergetics. 2010;2. Epub 2010/10/05. doi: 10.3389/fnene.2010.00026. PubMed PMID: 20890447; PubMed Central PMCID: PMCPMC2947927.

9. Fernandez-Moncada I, Robles-Maldonado D, Castro P, Alegria K, Epp R, Ruminot I, et al. Bidirectional astrocytic GLUT1 activation by elevated extracellular K(). Glia. 2021;69(4):1012–21. Epub 2020/12/06. doi: 10.1002/glia.23944. PubMed PMID: 33277953.

10. Ghergurovich JM, Garcia-Canaveras JC, Wang J, Schmidt E, Zhang Z, TeSlaa T, et al. A small molecule G6PD inhibitor reveals immune dependence on pentose phosphate pathway. Nat Chem Biol. 2020;16(7):731–9. Epub 2020/05/13. doi: 10.1038/s41589-020-0533-x. PubMed PMID: 32393898; PubMed Central PMCID: PMCPMC7311271.

11. Sotelo-Hitschfeld T, Niemeyer MI, Machler P, Ruminot I, Lerchundi R, Wyss MT, et al. Channel-mediated lactate release by K(+)-stimulated astrocytes. J Neurosci. 2015;35(10):4168–78. Epub 2015/03/13. doi: 10.1523/JNEUROSCI.5036-14.2015. PubMed PMID: 25762664; PubMed Central PMCID: PMCPMC6605297.

12. San Martin A, Arce-Molina R, Galaz A, Perez-Guerra G, Barros LF. Nanomolar nitric oxide concentrations quickly and reversibly modulate astrocytic energy metabolism. J Biol Chem. 2017;292(22):9432–8. Epub 2017/03/28. doi: 10.1074/jbc.M117.777243. PubMed PMID: 28341740; PubMed Central PMCID: PMCPMC5454122.

13. Almeida A, Moncada S, Bolanos JP. Nitric oxide switches on glycolysis through the AMP protein kinase and 6-phosphofructo-2-kinase pathway. Nat Cell Biol. 2004;6(1):45–51. Epub 2003/12/23. doi: 10.1038/ncb1080. PubMed PMID: 14688792.

14. Koberstein JN, Stewart ML, Smith CB, Tarasov AI, Ashcroft FM, Stork PJS, et al. Monitoring glycolytic dynamics in single cells using a fluorescent biosensor for fructose 1,6-bisphosphate. Proc Natl Acad Sci U S A. 2022;119(31):e2204407119. Epub 2022/07/27. doi: 10.1073/pnas.2204407119. PubMed PMID: 35881794; PubMed Central PMCID: PMCPMC9351453.

15. Vicente-Gutierrez C, Bonora N, Bobo-Jimenez V, Jimenez-Blasco D, Lopez-Fabuel I, Fernandez E, et al. Astrocytic mitochondrial ROS modulate brain metabolism and mouse behaviour. Nat Metab. 2019;1(2):201–11. Epub 2019/02/01. doi: 10.1038/s42255-018-0031-6. PubMed PMID: 32694785.

16. Taylor WM, Halperin ML. Stimulation of glucose transport in rat adipocytes by insulin, adenosine, nicotinic acid and hydrogen peroxide. Role of adenosine 3’:5’-cyclic monophosphate. Biochem J. 1979;178(2):381–9. Epub 1979/02/15. doi: 10.1042/bj1780381. PubMed PMID: 220963; PubMed Central PMCID: PMCPMC1186526.

17. Ben-Yoseph O, Boxer PA, Ross BD. Assessment of the role of the glutathione and pentose phosphate pathways in the protection of primary cerebrocortical cultures from oxidative stress. J Neurochem. 1996;66(6):2329–37. Epub 1996/06/01. doi: 10.1046/j.1471-4159.1996.66062329.x. PubMed PMID: 8632155.

18. Ben-Yoseph O, Boxer PA, Ross BD. Oxidative stress in the central nervous system: monitoring the metabolic response using the pentose phosphate pathway. Dev Neurosci. 1994;16(5-6):328–36. Epub 1994/01/01. doi: 10.1159/000112127. PubMed PMID: 7768213.

19. Kuehne A, Emmert H, Soehle J, Winnefeld M, Fischer F, Wenck H, et al. Acute Activation of Oxidative Pentose Phosphate Pathway as First-Line Response to Oxidative Stress in Human Skin Cells. Mol Cell. 2015;59(3):359–71. Epub 2015/07/21. doi: 10.1016/j.molcel.2015.06.017. PubMed PMID: 26190262.

20. Rodriguez-Rodriguez P, Fernandez E, Bolanos JP. Underestimation of the pentose-phosphate pathway in intact primary neurons as revealed by metabolic flux analysis. J Cereb Blood Flow Metab. 2013;33(12):1843–5. Epub 2013/09/26. doi: 10.1038/jcbfm.2013.168. PubMed PMID: 24064491; PubMed Central PMCID: PMCPMC3851909.

21. Bouzier-Sore AK, Bolanos JP. Uncertainties in pentose-phosphate pathway flux assessment underestimate its contribution to neuronal glucose consumption: relevance for neurodegeneration and aging. Front Aging Neurosci. 2015;7:89. Epub 2015/06/05. doi: 10.3389/fnagi.2015.00089. PubMed PMID: 26042035; PubMed Central PMCID: PMCPMC4436897.

